# SweetOrigins: Extracting Evolutionary Information from Glycans

**DOI:** 10.1101/2020.04.08.031948

**Authors:** Daniel Bojar, Rani K. Powers, Diogo M. Camacho, James J. Collins

## Abstract

Glycans, the most diverse biopolymer and crucial for many biological processes, are shaped by evolutionary pressures stemming in particular from host-pathogen interactions. While this positions glycans as being essential for understanding and targeting host-pathogen interactions, their considerable diversity and a lack of methods has hitherto stymied progress in leveraging their predictive potential. Here, we utilize a curated dataset of 12,674 glycans from 1,726 species to develop and apply machine learning methods to extract evolutionary information from glycans. Our deep learning-based language model SweetOrigins provides evolution-informed glycan representations that we utilize to discover and investigate motifs used for molecular mimicry-mediated immune evasion by commensals and pathogens. Novel glycan alignment methods enable us to identify and contextualize virulence-determining motifs in the capsular polysaccharide of *Staphylococcus aureus* and *Acinetobacter baumannii*. Further, we show that glycan-based phylogenetic trees contain most of the information present in traditional 16S rRNA-based phylogenies and improve on the differentiation of genetically closely related but phenotypically divergent species, such as *Bacillus cereus* and *Bacillus anthracis*. Leveraging the evolutionary information inherent in glycans with machine learning methodology is poised to provide further – critically needed – insights into host-pathogen interactions, sequence-to-function relationships, and the major influence of glycans on phenotypic plasticity.

---

In contrast to RNA and proteins, whose sequences can be elucidated from their DNA sequence, glycans are the only biological polymer that falls outside the rules of the central dogma of molecular biology. Glycans are present as modifications on all other biopolymers^1^, exerting varying effects on biomolecules, including stabilization and modulation of their functionality^2,3^. Apart from influencing the function of individual proteins, glycans are also crucial for cell-cell contact^4^ and mediate essential developmental processes^5^. Although glycans are synthesized by specific, DNA-encoded enzymes^6^, an individual glycan sequence is dependent on an intricate interplay between multiple enzymes and cellular conditions, providing glycans with important roles in phenotypic plasticity. Different glycosylation sites, even on the same protein, can exhibit considerable glycoform heterogeneity, depending on accessibility and glycosyltransferase kinetics, as has been shown for protein disulfide isomerase^7^. The glycan alphabet exceeds 1,000 monomers, allowing for an astronomical number of potential oligosaccharides built with different monosaccharides, lengths, connectivity, and branching.

Taken together, it can be appreciated that glycans possess great evolutionary plasticity, as sequences can be changed depending on conditions such as the level of extracellular metabolites^8^, without genetic mutations. Recently, Lauc et al. hypothesized that the plethora of available glycoforms and their plasticity facilitated the evolution of complex multicellular lifeforms^9^, reasoning that is well supported by the essential roles of glycans in developmental processes and cell-cell communication. As glycans make up the outermost layer of both eukaryotic and prokaryotic cells, any cross-kingdom interaction will necessarily involve glycans. Therefore, the prominent role of glycans in host-pathogen interactions^1^ has resulted in evolutionary pressures and opportunities on both sides. In short, as it has been put, nothing makes sense in glycobiology, except in the light of evolution^10^.

Given the rich evolutionary information contained in glycans, unique insights could be gained about pathogenicity and commensalism determinants as well as phylogenetics, the study of evolutionary relationships between taxonomic groups, from detailed analyses of glycan alignments and embeddings. Molecular mimicry of host glycans by both pathogens and commensals facilitates their immune evasion^11^, information that could be crucial to therapeutic efforts and glycoengineering probiotics. Established phylogenetic approaches relying on the bacterial 16S rRNA gene^12^ are limited in resolution by the requirement of 16S rRNA variation between taxa, while recent studies have demonstrated that capsular polysaccharides in fungi can be used to resolve and strengthen DNA-based phylogenetic trees^13^. This not only emphasizes the complementary evolutionary information present in glycans but also strengthens our hypothesis of using glycan alignments and embeddings for phylogenetic inferences.

We recently established a deep learning-based language model for glycans, SweetTalk^14^. Language models are well suited for leveraging patterns and implicit structure in biological sequences such as nucleic acids^15^ and proteins^16^. Analogously, we trained SweetTalk by extracting characteristic trisaccharides from glycans, referred to as “glycowords”, and using their sequence context to learn the language of glycans. This approach to biological sequences not only enables learning a representation of a molecule but also offers the potential for transfer learning − by training a deep neural network on raw sequences, it is possible to transfer fundamental principles inherent in any given biopolymer and train a classifier on a smaller dataset of sequences associated with labels or metadata, such as glycan linkage or immunogenicity to humans.

Here, we introduce SweetOrigins, a language model-based approach that enables one to extract evolutionary information from glycan sequences by training multiclass classifiers with high accuracy for every taxonomic level. To achieve this objective, we curated a comprehensive dataset comprising species-specific evolutionary information for 12,674 glycans that we used for motif analyses as well as biomining. These data, together with our previously described glycan dataset^14^, equipped us with a large database, GlycoBase, that is amenable to programmatic access and integration into deep learning pipelines. We leveraged the evolutionary information gained by SweetOrigins to identify and analyze glycan motifs used for molecular mimicry-mediated immune evasion by commensals and pathogens, such as the Lewis blood antigens. Utilizing glycan alignments, we also discovered and contextualized motifs relevant for pathogenicity, with the example of the enterobacterial common antigen (ECA) and capsular polysaccharides of *Staphylococcus aureus* and *Acinetobacter baumannii*. Additionally, we used both glycan alignments and evolution-informed glycan representations to construct phylogenetic trees based on glycan sequences alone. Glycan-based phylogenies recovered most of the information contained in 16S rRNA-based phylogenies and outperformed DNA phylogenies for differentiating genetically similar species with diverging phenotypes, such as *Bacillus cereus* and *Bacillus anthracis*. As we will describe below, SweetOrigins and our glycan-based phylogenies offer a powerful approach for including phenotypic plasticity into phylogeny and discovering as well as contextualizing functional motifs involved in pathogenicity.

## Results

### Using a large species-specific glycan dataset for motif analysis and biomining

Leveraging information from UniCarbKB, the Carbohydrate Structure Database (CSDB), and targeted literature searches equipped us with 12,674 glycan sequences and their species association (see Methods; Supplementary Table 1), containing thousands of diverse glycans absent from our previously reported sequence dataset^14^. Combining both datasets (Fig. 1a, Supplementary Table 2) nearly doubled the number of unique glycoletters (monosaccharides or bonds) present in glycans from 602 to 1,027 and those of glycowords (i.e., extracted trisaccharides) from 8,843 to 15,155 unique glycowords, which can be further increased to 19,866 glycowords by considering isomorphic glycans (Supplementary Fig. 1). Not only was the vocabulary of utilized glycowords larger in the combined dataset, the increase of observed glycoletters now allowed for the theoretical formation of more than one trillion possible glycowords. This, together with the fact that we only observe ∼0.0000013% of these glycowords in our dataset, indicates that there is substantially more diversity in unknown glycans. Equipped with this extensive dataset, we set out to analyze the characteristic local structural environments of glycoletters (Supplementary Fig. 2). With the monosaccharide fucose as an example, we observed N-acetylglucosamine (GlcNAc) and galactose (Gal) as typical connected monosaccharides (Fig. 1b), concordant with the fucosyltransferase substrate specificities annotated in glycosyltransferase family 10^17^. Additionally, we ascertained that fucose had an overall higher likelihood of occurring in the main branch of a glycan rather than its side branches. This information, together with the sharp drop-off in observed binding after galactose, could be useful in future efforts to reconstruct glycan sequences based on maximum likelihood inferences, as most glycoletter combinations will lack corresponding glycosyltransferases and therefore be highly unlikely, as also indicated by the clear separation of glycoword clusters in embedding space (Supplementary Fig. 3-4).

**Table 1.**
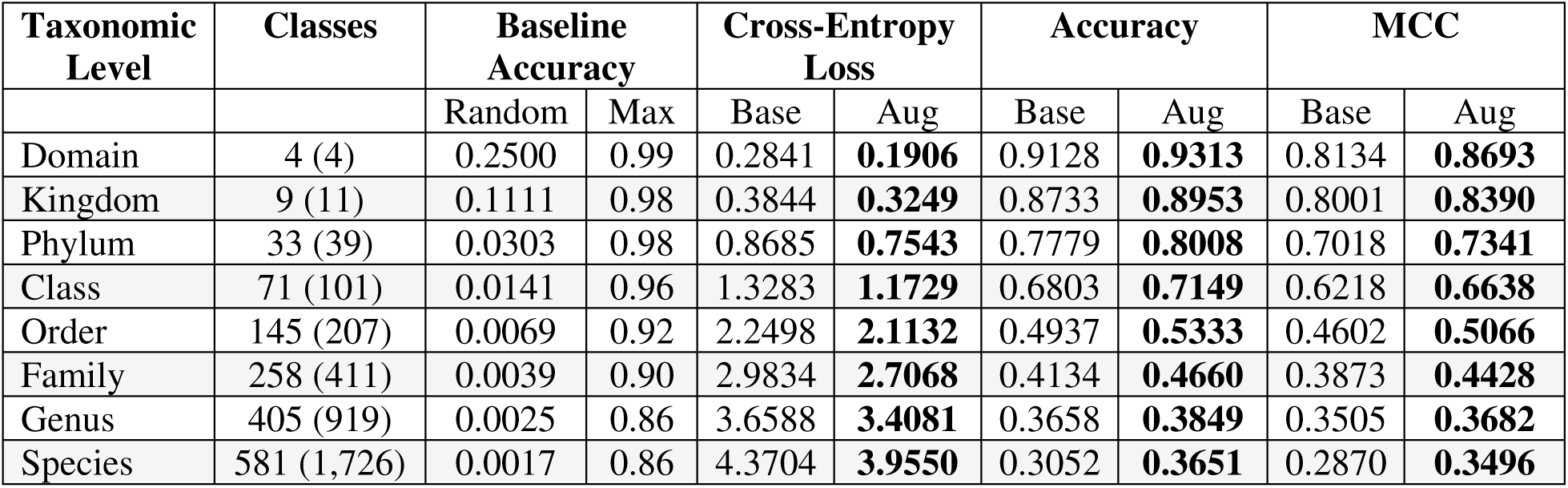
Metrics of trained SweetOrigins models. For every taxonomic level, glycans were split into training (80%) and validation (20%) sets as described in the Methods section. Taxonomic groups with fewer than five unique glycans were not used for model training or validation. Number of classes indicates the number of included taxonomic groups, while the full number of taxonomic groups in our dataset is given in parentheses. Models were trained with the standard set of glycans (Base) or after data augmentation (Aug). As a baseline accuracy, a random prediction of classes was used for each model. Max indicates the maximum theoretically possible accuracy given the glycan sequences that are shared across taxonomic groups. Cross-entropy loss, accuracy, and Matthew’s correlation coefficient (MCC) of the trained model on a separate validation set are given for each taxonomic level. For each metric and taxonomic level, the superior value is indicated in bold.

**Fig. 1.**
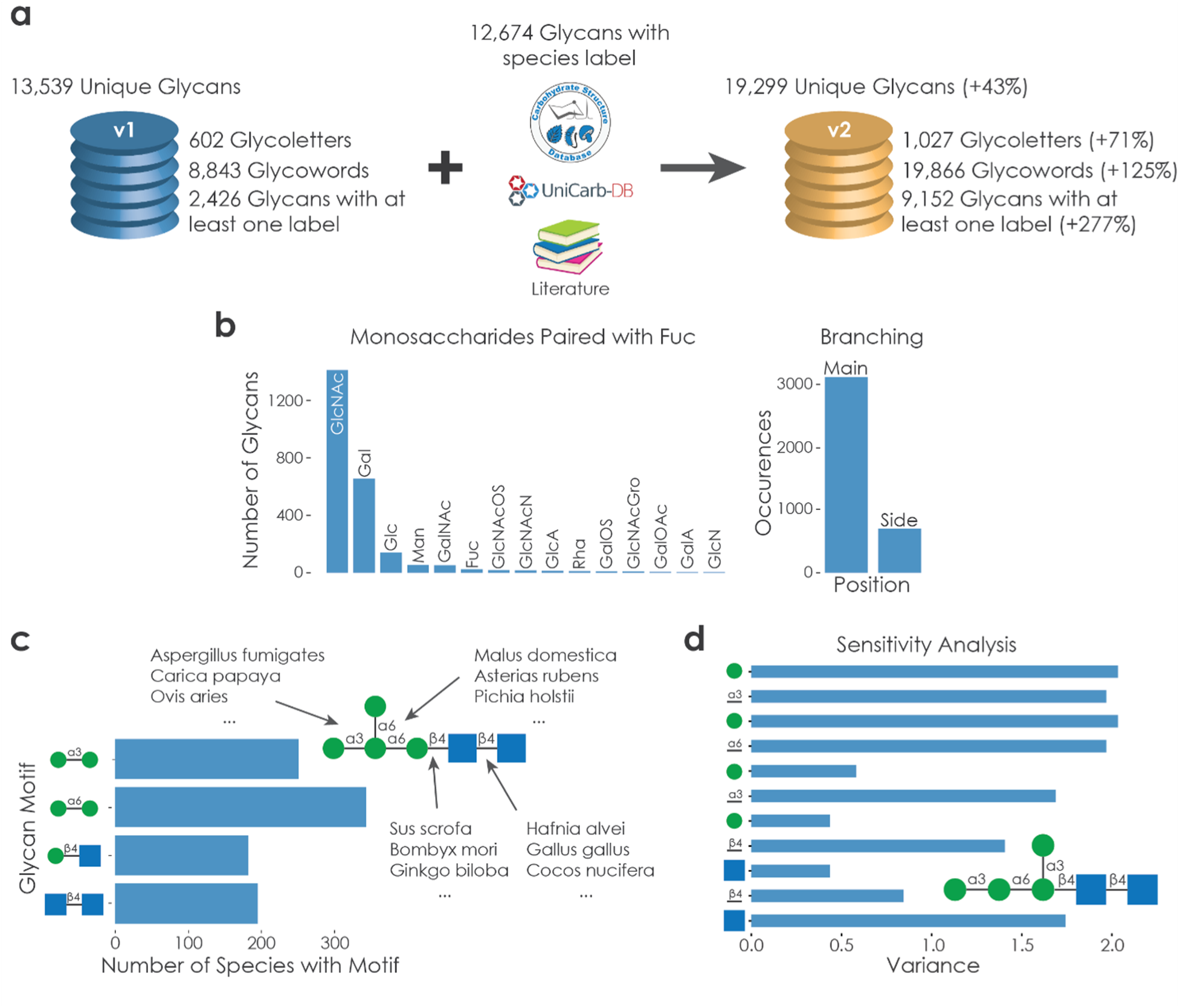
Using a large glycan dataset for analyzing glycan substructures and predicting biomining targets. **a**, Building a large dataset of labeled glycan sequences. A previously reported glycan database^14^ (v1) was merged with glycan sequences including species information from CSDB (carbohydrate structure database), UniCarbKB, and the peer-reviewed scientific literature. This resulted in an expanded database (v2) that not only contained additional glycan sequences but also more associated metadata, such as species or linkage information. **b**, Analyzing the local structural context of glycoletters. As an example, we ordered the frequency of all monosaccharides following fucose in the glycans present in our dataset, highlighting its characteristic local structural context together with its likely position in the glycan structure (main versus side branch). **c**, Utilizing link libraries for glycan biomining hypothesis generation. For all glycans in our database with species information, we constructed a library of disaccharide motifs that are present in any species to generate leads for glycosyltransferase biomining. **d**, Glycan sensitivity analysis. Given a glycan sequence, we exhaustively performed *in silico* modification, only retaining substitutions that resulted in observed glycowords. The number of allowed substitutions at any position was used to calculate sequence variance by normalizing to the total amount of observed monosaccharides or bonds. Glycans are drawn in accordance with the symbol nomenclature for glycans (SNFG).

**Fig. 2.**
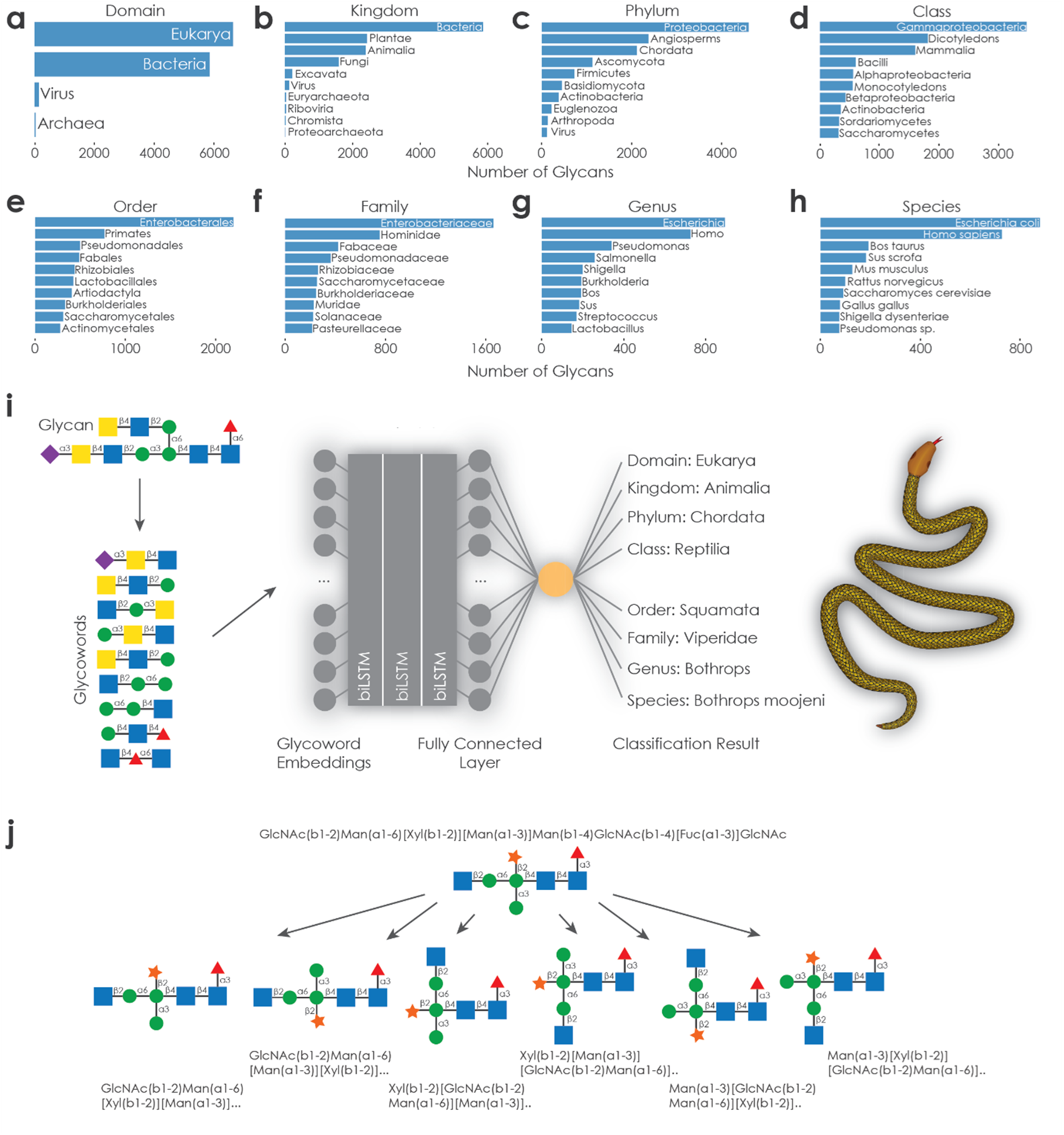
Glycan species distribution and methods to extract evolutionary information from glycan sequences. **a-h**, For all glycans labeled with species information, we determined up to the 10 most abundant classes of the taxonomic levels of domain (**a**), kingdom (**b**), phylum (**c**), class (**d**), order (**e**), family (**f**), genus (**g**), and species (**h**). **i**, Exemplary schematic of SweetOrigins to extract evolutionary information from glycan sequences. Any given glycan sequence is featurized into a set of glycowords. This list of glycowords is used as input for a trained SweetOrigins model, consisting of a three-layered, bidirectional recurrent neural network with a preceding embedding layer and a subsequent fully connected layer, predicting the taxonomic class (depending on the specific model). Trained models range from the domain level down to the species level. **j**, Glycan data augmentation strategy. As described in the Methods section, different bracket notations describing the same glycan can be generated by alternating double branches as well as replacing side branches with main branches to increase model robustness.

**Fig. 3.**
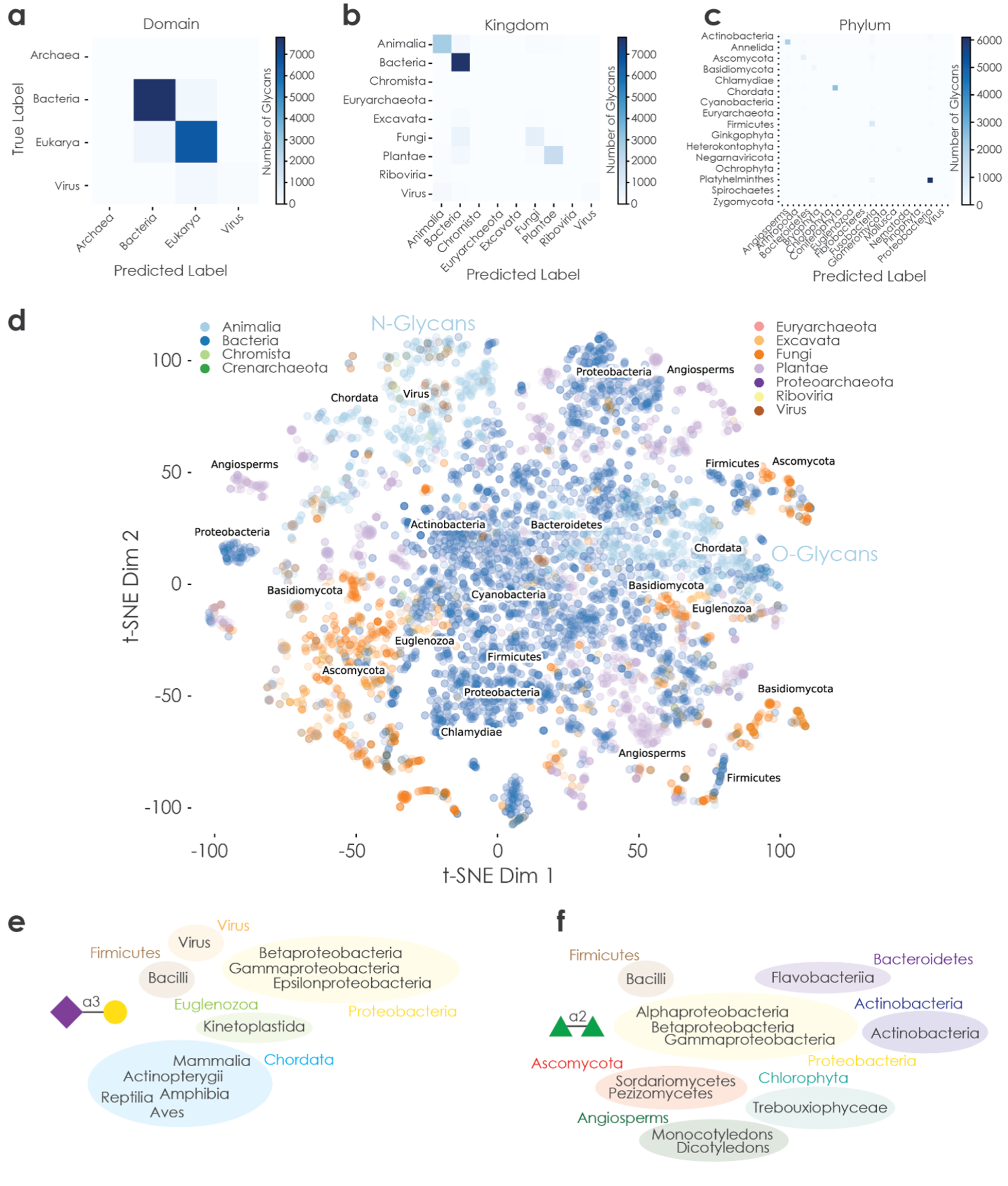
Machine learning-based classifiers predict the taxonomic origin of glycans. Confusion matrices of the domain-level (**a**), kingdom-level (**b**), and phylum-level (**c**) SweetOrigins models. For all glycans of taxonomic groups at the respective level with at least five glycans, true taxonomic labels were plotted against the labels predicted by the trained models, showcasing correct predictions on the diagonal. For the phylum-level confusion matrix, labels are shown alternatingly on each axis. **d**, Glycans in evolutionary embedding space separate according to taxonomic groups. For all glycans with species information, an embedding was formed from the trained embedding layer of the phylum-level SweetOrigins model, shown via t-distributed stochastic neighbor embedding (t-SNE). This embedding was colored by the kingdom from which a given glycan stemmed. Then, clusters of glycans belonging to representative phyla were labeled with that phylum. Clusters of vertebrate glycans are labeled to indicate whether they contain N- or O-linked glycans. **e**, Evolutionary tracking of disaccharide motifs. For a given disaccharide motif, we searched all glycans with species information, collected all taxonomic classes that contained glycans with the respective motif, and colored them by their phyla.

Cataloguing all disaccharide motifs present in species in our glycan database yielded a list of species that can synthesize a given glycan bond and that could be targeted for enzyme biomining for glycoengineering or related purposes (Fig. 1c, Supplementary Fig. 5). This species-based analysis revealed that some disaccharides, such as Man(α1-6)Man, are present across hundreds of species, whereas others, such as Man(α1-2)Fuc, are restricted to only a few species, making it more challenging to find suitable enzymes for glycoengineering and stressing the importance of our biomining approach. Importantly, identifying targets for biomining based on reaction products represents an orthogonal approach to current methods of searching for enzyme DNA sequences homologous to known glycosyltransferases. Analogous to protein engineering, knowing which regions of a biopolymer to modify is crucial. Thus, we performed *in silico* modification of glycan sequences (see Methods) to assess how conserved a given region of a glycan was. These sensitivity analyses revealed positions in glycans that are near-constant, together with regions such as the non-reducing end that show high variance (Fig. 1d, Supplementary Fig. 6). As conserved sequences are more likely to be functionally relevant, future glycoengineering efforts could profit from focusing on these variable regions and avoid changes to conserved, potentially functionally relevant glycan regions.

To facilitate further advances in glycobiology, we created GlycoBase, a comprehensive glycan database with metadata and analytical tools based on this work (Supplementary Fig. 7a, Supplementary Table 3, https://wyss.shinyapps.io/glycobase/). GlycoBase offers readily accessible glycan data, explorable glycan embeddings, and many of the methods developed here as tools, such as the local structural context of any glycoletter (Supplementary Fig. 7b). We envision that this resource will help accelerate progress in the development and application of machine learning methods to glycobiology and glycoengineering.

### Extracting evolutionary information from glycan sequences with SweetOrigins

Our labeled dataset contained glycans from 1,726 species from 39 taxonomic phyla, representing a comprehensive snapshot of currently known species-specific glycans (Fig. 2a-h). Taking into account the hierarchical nature of taxonomy, we developed SweetOrigins, a multiclass classifier trained on glycan sequences with the same base model architecture on every taxonomic level (Fig. 2i). By initializing SweetOrigins with glycoword embeddings learned by SweetTalk, trained on a larger, unlabeled set of glycans, our resulting models achieved high levels of validation accuracy and Matthew’s correlation coefficients (MCCs) on all taxonomic levels (Table 1). We further employed data augmentation by capitalizing on the notion of glycans as graphs and formed a set of isomorphic graphs comprising slightly different lists of glycowords (Fig. 2j). This led to model performance improvements at every classification level, with absolute accuracy increases of up to six percent, and could make glycan models more robust in general, analogous to data augmentation in image classification^18^. SweetOrigins presents an important benchmark for future glycobiology-focused machine learning efforts and its success clearly demonstrates the abundant evolutionary information contained in glycans that can be leveraged for predictive purposes.

While SweetOrigins demonstrates excellent classification performance, viral glycans are frequently misclassified (Fig. 3a). Given that viruses predominantly derive their glycans from their hosts, viral glycan sequences from human immunodeficiency virus (HIV) and the current SARS-CoV-2 pandemic, respectively, typically match those of their host organisms (Supplementary Fig. 8). In general, our predictions were robust and any misclassifications occurred among closely related taxonomic groups, including some fungal glycans falsely predicted to be derived from bacteria on the kingdom level (Fig. 3b) and Firmicutes glycans mislabeled as those of Proteobacteria on the phylum level (Fig. 3c). As glycan embeddings can be constructed from models trained on every taxonomic level, we quantified embedding distances across models from different taxonomic levels (Supplementary Fig. 9). This analysis revealed that the embeddings for shorter glycans changed more dramatically, potentially due to their lower expressiveness, as longer glycans comprised more glycowords and therefore provided more robust predictions.

Glycan embeddings derived from the trained SweetOrigins models illustrated evolutionary clusters reminiscent of the taxonomic groups (Fig. 3d). Additionally, vertebrate glycans formed separate clusters for their N- and O-linked glycans, respectively, indicating a rich representation learned by SweetOrigins. Intriguingly, the cluster containing O-linked glycans, predominantly present on mucosal surfaces, partially overlapped with the bacterial glycan cluster. Further, the closest bacterial phyla, Bacteroidetes, Firmicutes, and Proteobacteria, are common constituents of the vertebrate microbiome^19^.

We also tracked glycan motifs in the trained evolutionary glycan embeddings. We found, for example, that NeuNAc(α2-3)Gal, the binding motif of the innate immunity complement control protein Factor H^20^, was present in parasitic Euglenozoa causing leishmaniasis and Chagas disease as well as both vertebrate and bacterial classes (Fig. 3e), with an enrichment for bacterial classes close to vertebrate O-linked glycans in embedding space (Supplementary Fig. 10a). The absence of this motif in plants and fungi supports its separate and independent emergence in vertebrates and bacteria^21^, especially commensals and pathogens. While this motif was also found in protozoans, trypanosomes have been shown to be capable of acquiring sialic acid from host glycans to decorate their own glycans^22^. Our tracking process also demonstrated the widespread abundance of Rha(α1-2)Rha in bacteria, fungi, and plants (Fig. 3f), while it was absent from any vertebrate classes. Combined with a tight clustering of glycans containing this motif in embedding space (Supplementary Fig. 10b), this hints at a common evolutionary origin of this motif.

We also employed our evolutionary tracking approach for probing functional relationships of glycan motifs. The disaccharide motif Fuc(α1-2)Gal is known for its role in cancer^23^ and is essential for the formation of Lewis blood antigens. Evolutionary tracking confirmed previous findings from Ma et al.^24^, demonstrating the presence of Fuc(α1-2)Gal in glycans of vertebrates, nematodes, Basidiomycota, Proteobacteria, and others (Supplementary Fig 11). In the evolutionary embeddings, Fuc(α1-2)Gal is enriched in the vertebrate O-linked glycans and surrounding bacterial glycans. Indeed, bacteria such as *Helicobacter pylori* synthesize blood group antigens to escape the human immune system by molecular mimicry^25^. Mice lacking the critical galactoside 2-alpha-L-fucosyltransferase 2 (FUT2), and thus lacking the corresponding Lewis blood antigen, are largely resistant to *H. pylori* infection^26^. Other instances of molecular mimicry of host glycans^11^ suggest a broader role of this phenomenon and point to an important opportunity for the approach presented here to identify mimicry motifs as well as the commensal and pathogens exhibiting them. Notably, we identified many additional pathogens or parasites, such as the nematodes *Dirofilaria immitis* and *Toxocara canis*, with Fuc(α1-2)Gal in their glycans, potentially as molecular mimicry (Supplementary Fig. 11).

Even though SweetOrigins classifiers were only trained up to the species level, we demonstrated that even strain-level information could be extracted from glycans by using their learned representation from the species-level SweetOrigins model (Supplementary Fig. 12). For example, using the 1,010 glycan sequences from the model organism *E. coli* with strain-level annotation, we readily identified clusters enriched for several strains, such as the serotypes O8/O9, characterized by a special polymannose O-antigen^27^, or the K-12 strain popular in molecular biology research.

### Glycan alignments contextualize pathogenic motifs in *S. aureus* and *A. baumannii*

A common method of analyzing motifs in biological sequences that capitalizes on evolutionary information is the use of alignments. We thus developed methods for gapped, pairwise alignments of glycan sequences, assisted by a substitution matrix analogous to the BLOSUM matrices utilized in protein alignments (which we termed GLYSUM; Supplementary Table 4). Based on our expansive dataset comprising 1,027 monosaccharides or bonds, alignments of highly diverse bacterial glycans are also possible. To demonstrate this, we aligned the serotype 5 capsular polysaccharide of the clinically relevant pathogen *S. aureus*, that is known to increase bacterial virulence^28^, against our entire dataset (Fig. 4a). Notably, the best alignments were achieved with the enterobacterial common antigen (ECA), conserved in the Enterobacteriaceae family, that has been shown to be important for virulence^29^ and outer membrane permeability^30^. This alignment result is supported by experiments demonstrating that ECA deficiency in *E. coli* can be complemented by the expression of enzymes from serotype 5 *S. aureus*^31^. Given that the capsular polysaccharides of *S. aureus* mediate its evasion of the immune system^32^, any insight into shared epitopes with the commensals from the Enterobacteriaceae family is potentially valuable for understanding and combating its pathogenicity.

**Fig. 4.**
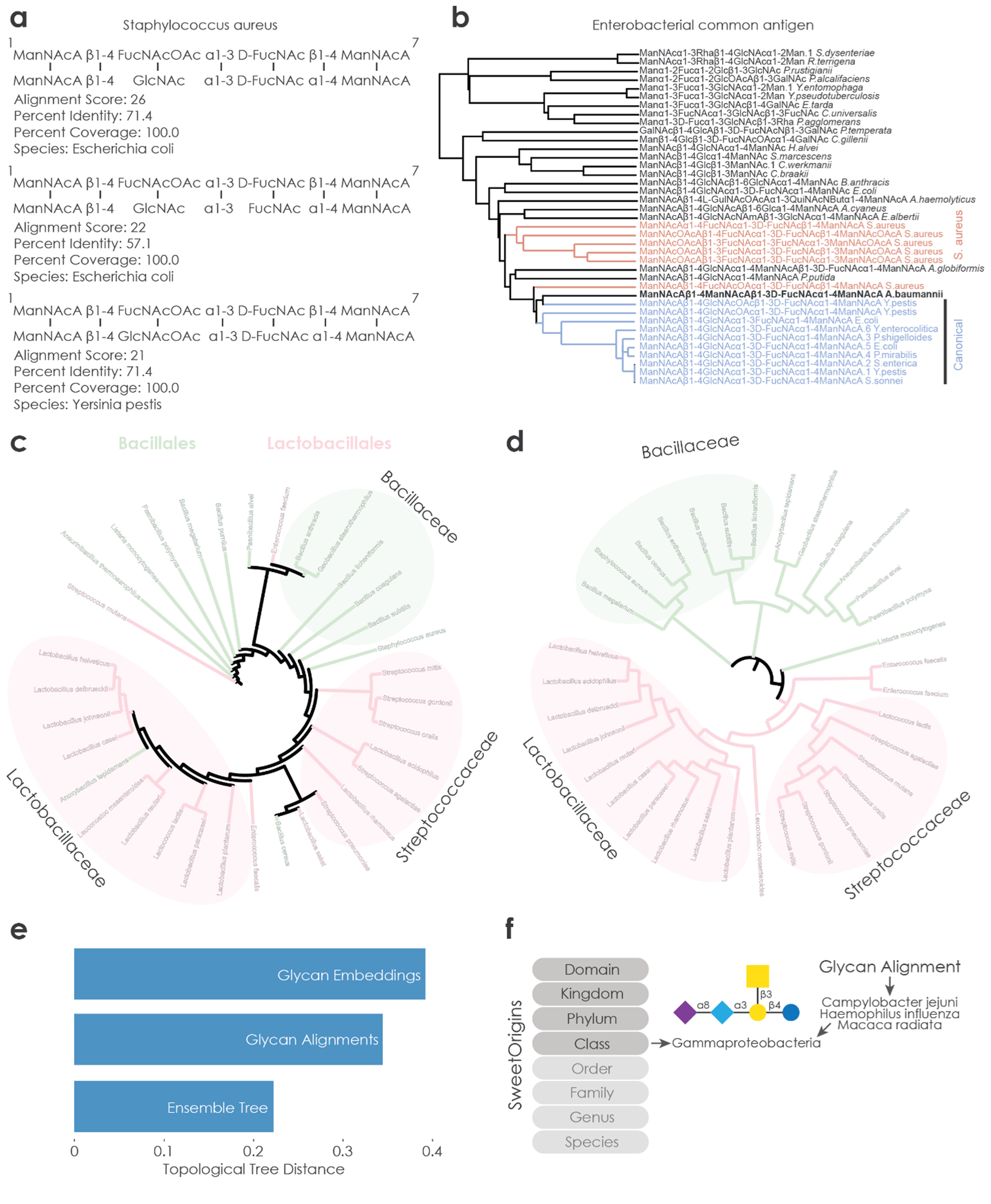
A glycan-based phylogeny to gain an improved understanding of evolution. **a**, Glycan alignments using serotype 5 capsular polysaccharide of *S. aureus*. The repeating unit of the glycan was aligned against the whole annotated database and the best three alignments are shown with species information. **b**, Enterobacterial common antigen (ECA) and ECA-like glycans. We aligned the canonical ECA sequence against our entire dataset and curated ECA-like sequences from the best 50 alignments. A dendrogram resulting from alignment distances is shown together with ECA-like sequences and their species of origin. **c**, Phylogenetic ensemble tree of Bacilli based on glycan alignments and embeddings. For all species in the taxonomic class Bacilli with at least five glycans in our dataset, we averaged their embedding as well as alignment distances and used hierarchical clustering to construct a phylogenetic tree, visualized by Interactive Tree of Life v5.5^33^. Species are colored according to their taxonomic order and clusters attributable to a taxonomic order are colored as well. **d**, Phylogenetic tree of Bacilli using 16S rRNA alignment. Based on multiple sequence alignments, the phylogenetic tree for Bacilli in our dataset, derived from neighbor joining, is shown and colored according to (c). **e**, Topological distance between phylogenetic trees of Bacilli. Branch length-aware topological distance, as implemented in the R package ape^34^ (version 5.3), was calculated between the 16S rRNA-based Bacilli phylogeny and all glycan-based trees reported in this work. **f**, Using SweetOrigins and glycan alignments to predict evolutionary origin of unlabeled glycans. Glycan sequences are used as inputs for trained SweetOrigins models as well as for exhaustive pairwise alignment. The first instance of concordance between both modalities is used as a taxonomic prediction of the origin of any given glycan.

Aligning the canonical ECA with our glycan dataset allowed for the compilation of a list of ECA-like glycan sequences and their alignment distances that we used to construct a dendrogram (Fig. 4b). While most of the *S. aureus*-derived ECA-like sequences formed a separate cluster, the type 5 capsular polysaccharide was firmly located in the cluster containing the canonical ECA sequences, potentially explaining its elevated virulence. We further observed an ECA-like glycan for *Acinetobacter baumannii* (Fig. 4b, bold), one of the most devastating hospital-acquired pathogens, in the same cluster dominated by canonical ECA sequences. The capsular polysaccharide of *A. baumannii* has been identified to be involved in antibiotic resistance and virulence^35^, providing a link to the functions of canonical ECA. For other pathogens, such as *Haemophilus ducreyi*, the expression of a gene cluster synthesizing a putative, ECA-like glycan has also been linked to increased virulence^36^, demonstrating the importance of this motif. The genera Staphylococcus, Acinetobacter, and Haemophilus are not part of the Enterobacteriaceae family typically associated with the ECA, clearly demonstrating the relevance of our glycan alignments for identifying and contextualizing motifs important for pathogenicity, such as their ECA-like glycans.

### Establishing a glycan-based phylogeny to distinguish phenotypically divergent species

The evolutionary information in glycans, in addition to its utility for predicting the origin of a given glycan of interest, could also serve as a basis for a glycan-based phylogeny, describing the relationships between taxa. Phylogenetic trees commonly utilize a distance-based measure to determine branching. Accordingly, we used the distance between glycan embeddings derived from SweetOrigins as a measure of evolutionary distance. Glycans, in contrast to genes, are too diverse to identify sequences that are shared by a large swath of the evolutionary spectrum and that could be used as phylogenetic markers. Thus, we calculated the average glycan embedding for every species as a characteristic snapshot of its glycome. In addition to providing a basis for an evolutionary distance, this approach allowed us to disentangle viral glycans from those of their hosts, which were indistinguishable on the level of individual glycans. While every single viral glycan is host-derived, the sum of viral glycans only presents a subset of the host glycome and therefore results in different average glycome embeddings (Supplementary Fig. 13).

Performing hierarchical clustering on the resulting distance matrix enabled us to construct a glycan-based evolutionary tree for the well-studied taxonomic class of Bacilli in the phylum Firmicutes (Supplementary Fig. 14a), containing several important commensals and pathogens; this allowed us to observe a partial disentanglement of the constituting taxonomic orders Lactobacillales and Bacillales.

To facilitate comparison with established alignment-based phylogenies, we computed pairwise alignments between all glycans of two species. This yielded an average distance between them, allowing us to calculate a distance matrix as well as the corresponding phylogenetic tree (Supplementary Fig. 14b). Evolutionary trees based on glycan alignments exhibited more clearly recognizable evolutionary clades than corresponding trees based on glycan embeddings. Reasoning that our evolutionary embeddings and glycan alignments captured different aspects of glycan biology, we then formed an ensemble tree by combining phylogenetic trees derived from embeddings and alignments (Fig. 4c). The resulting tree indeed exhibited an improved clustering of Lactobacillaceae and Streptococcaceae for Lactobacillales and Bacillaceae for Bacillales, respectively, on the taxonomic level of family.

Next, we sought to compare our glycan-based phylogenetic tree to the common phylogenetic approach applied to bacteria, that relies on the alignment of the 16S ribosomal RNA as a phylogenetic marker gene (Fig. 4d). The notable bisection of Lactobacillales and Bacillales by this phylogenetic tree clearly supported our ensemble tree based on glycans, demonstrating that a purely glycan-based phylogeny is not only feasible but also highly informative. By quantifying the topological difference between the 16S rRNA-based phylogeny and glycan-based phylogenetic trees, we could again establish the superiority of the ensemble approach over using either glycan embeddings or alignments separately (Fig. 4e).

Interestingly, the common food and soil microbe *Bacillus cereus* was indistinguishable from the causative agent of anthrax, *Bacillus anthracis*, on the 16S rRNA level, yet was readily differentiated by the glycan-based phylogeny. Based on DNA sequencing, it has even been suggested that *B. cereus, B. anthracis*, and the agriculturally relevant *B. thuringiensis*, should be aggregated into a single species^37^. While these bacteria only differ by their acquired plasmids, they differ dramatically in their phenotype. This difference is reflected in their glycans, as one of the differences between these bacteria is the expression of additional glycosyltransferases by *B. anthracis*^38^. As such, seemingly minor differences at the genetic level are translated into large differences at the glycan level, reinforcing the need for a glycan-based phylogeny as a differentiator of species such as *B. cereus* and *B. anthracis*. As seen with molecular mimicry, glycans can be an essential constituent of pathogenicity and therefore are important for the phenotypic distinction between closely related species, underlining the importance of the glycan-based phylogeny approach presented here.

The species classifiers and glycan alignment tools developed in this work proved useful both for investigating evolutionary relationships and for inferring properties of existing glycans. To demonstrate this, we used the 10,333 unlabeled glycans in our dataset and utilized both the trained SweetOrigins models as well as glycan-based alignments to infer the species (or higher taxonomic group in case of uncertainty) any glycan most likely stemmed from (Fig. 4f, Supplementary Table 5). This analysis produced thousands of testable predictions, highlighting its potential for future efforts to catalog the diversity and understand the function of glycans in different species. For several predicted glycans, we engaged in targeted literature searches to validate the predictions made by SweetOrigins (Supplementary Fig. 15), confirming our approach and offering a tool for researchers aiming to identify the origins of glycans in complex mixtures.

## Discussion

Despite their incredible diversity and abundance, glycans have until now been hardly utilized as a source of evolutionary information on a larger scale. Speculations regarding the overarching evolutionary role of glycans notwithstanding, current phylogenetic approaches disregard information from this critical biopolymer existing outside the central dogma of biology. Here, we presented a collection of deep learning and bioinformatics strategies for how to extract information from glycan sequences. The high accuracy exhibited by our SweetOrigins models clearly demonstrated that glycans can be used to distinguish closely related taxonomic groups, including critical classification at the species level. Our results provide benchmarks to facilitate comparison of machine learning approaches extracting information from glycan sequences and could spur further advances in glycan-based machine learning, both methodologically as well as insight-deriving, transforming glycobiology into a domain that is both predictive as well as engineerable.

The glycan alignment methods we developed here hold promise by advancing phylogenetic approaches via incorporating phenotypic information and could also readily be used to identify glycans from complex mixtures, such as those derived from soil or clinical samples. In contrast to metagenomics, glycan alignments from these samples could provide immediate mechanistic insight into phenotypic characteristics. For glycans shared across organisms with variations, glycan alignments could identify and analyze conserved areas, similar to the broad applications of DNA and protein alignments, providing functional information and opportunities for glycoengineering. As with all other modules presented here, glycan alignments will become increasingly more powerful as glycan databases with species information expand by continuing research efforts. Along these lines, we developed a graphical user interface in GlycoBase to perform glycan alignments, similar to the popular BLAST alignment tools.

Multiple avenues presented in this work − evolutionary tracking, glycan alignments, glycan classification are suitable for connecting glycan function with sequence patterns or substructures, and are available on GlycoBase. As glycans are a major facilitator of phenotypic plasticity^1^, this endeavor is crucial for differentiating pathogens from closely related species, elucidating their pathological determinants, and investigating the differing phenotypes of the same organism under different conditions. The methods developed here enable rapid discovery, understanding, and utilization of functionally relevant glycan motifs, especially in the context of pathogens. Glycan motifs contributing to virulence, such as the capsular polysaccharides analyzed here, could thus be of broad relevance for diagnostics, vaccines, and targeted therapies.

The focus of the scientific community on a subset of (model) organisms, known as taxonomic bias, and the relative scarcity of viral and archaeal glycans in the current academic literature offer future opportunity for improvement, especially as glycosylation is hypothesized to have initially evolved in archaea^39^ and glycan diversity is expected to be very high in that domain of life. As research on archaeal glycans progresses, SweetOrigins and GlycoBase can be readily updated and will enable an even more comprehensive investigation of the evolutionary path and diversity of glycans. Further advances in glycobiology will eventually allow for precise evolutionary classification at the strain level, in addition to the species level.

A common theme of the motifs analyzed in this work points to the broader principle of molecular mimicry for both commensals and pathogens to evade the host immune system. Equipping microorganisms with identified immune evasion glyco-motifs, aided by immunogenicity-predicting deep learning models^14^, could enable the creation of synthetic biology-inspired, gut-resident sentinel bacteria^40^ with prolonged viability and exceptionally low immunogenicity. Evolutionary tracking and glycan alignments could thus prove important for the identification of organisms engaging in molecular mimicry and pinpointing their evolved strategies for imitating host glycans, potentially paving the way for targeted therapeutics.

## Methods

### Dataset

To create a comprehensive glycan dataset annotated with species labels, we manually curated 12,674 glycan sequences from three sources: UniCarbKB^41^, the Carbohydrate Structure Database (CSDB)^42^, and the peer-reviewed scientific literature. From UniCarbKB, we compiled all glycans with species information, a length of at least three monosaccharides, and a working link to PubChem. We further complemented and extended this list by gathering all glycans deposited in the Carbohydrate Structure Database (CSDB) up to December 2019 with a length of at least three monosaccharides. For species with more than 15 strains available on CSDB, only glycans from the first 15 strains were recorded to prevent taxonomic bias. For the model organism *Escherichia coli*, all available glycan sequences were recorded to facilitate a strain-based analysis. Finally, we performed additional literature searches, predominantly adding viral and archaeal glycans (Supplementary Table 6), which are underrepresented in the other databases. We revised and completed the annotations for all species’ taxonomic characterization (species, genus, family, order, class, phylum, kingdom, domain) based on the NCBI Taxonomy Browser. In total, the dataset contained sequences from 1,726 different species from a range of 39 taxonomic phyla.

To the best of our knowledge, this database represents the most comprehensive and current resource of glycans and their species information to date (Supplementary Table 1).

To enable transfer learning, we merged this dataset with a set of glycans we previously reported^14^ (here referred to as GlycoBase v1), resulting in an augmented database of 19,299 unique glycan sequences (Supplementary Table 2; GlycoBase v2). These glycans were paired with an ID to allow for a relational database linking all available information (linkage type, species information, human immunogenicity, etc.) to a glycan sequence (Supplementary Table 3).

### Data processing

Glycan sequences were processed as described previously^14^. Briefly, we removed dangling bonds and position-specific information of monosaccharide modifications. Then, we harmonized capitalization and, in the case of glycan repeat structures, appended the first monosaccharide to their end to capture more sequence context. Additional steps to exclude duplicated glycans included strict ordering of multiple branches with equal lengths by ascending connection to the main branch (e.g., branch ending in ‘α1-2’ before branch ending in ‘β1-4’). For outermost branches, the longest branch was defined as the main branch. Additionally observed monosaccharide modifications necessitated an expansion of the hierarchy to avoid mislabeling: NAc > OAc > NGc > OGc > NS > OS > NP > OP > NAm > OAm > NBut > OBut > NProp > OProp > NMe > OMe > CMe > NFo > OFo > OPPEtn > OPEtn > OEtn > A > N > SH > OPCho > OPyr > OVac > OPam > OEtg > OFer > OSin > OAep > OCoum > ODco > OLau > OSte > OOle > OBz > OCin > OAch > OMal > OMar > OOrn > rest.

Data processing for model training, as previously described^14^, included featurization of glycan sequences into pure characters (e.g., ‘G’, ‘a’, ‘l’), glycoletters (e.g., ‘Gal’), as well as glycowords (three monosaccharides connected by two bonds). The conversion of a glycan sequence into glycowords resulted in a list of partially overlapping glycowords, comprehensively representing the structural contexts in a given glycan.

All abbreviations for glycan nomenclature in this work can be found in Supplementary Table 7.

### Analyzing links in glycan sequences

To determine typical local structural contexts of monosaccharides and bonds, we quantified the frequency of a given monosaccharide co-occurring with any other monosaccharide in our extensive database of unique glycans. Additionally, we also compared the relative frequencies of a particular monosaccharide being observed in the glycan main branch versus a side branch in of our database.

We also enumerated all observed disaccharides (e.g., Man(α1-6)Man) by their presence in each species, in order to identify species likely to express the enzymes for catalyzing particular bonds.

### Glycan *in silico* modification

We performed *in silico* modification of glycans by replacing monosaccharides and/or bonds with other observed monosaccharides/bonds, as described in prior work^14^. For the current study, we used exhaustive modification, replacing glycoletters with all possible glycoletters while only retaining modified glycans comprising previously observed glycowords. This ensured physiological relevance, given the extreme sparsity of observed glycan sequences compared to the theoretical number of possibilities.

The number of alternative glycoletters allowed at any given position in a glycan was used as a proxy for how conserved this region was. Under this assumption, sensitivity analyses of glycans can be performed to identify variable and conserved regions. For these analyses, all unique substitutions occurring at a given position were assessed and divided by the log-transformed number of total possible monosaccharides or bonds, depending on the position, to yield a measure of variance.

### Glycan alignment

Global sequence alignment of glycans was implemented according to the Needleman-Wunsch algorithm by adapting the Python Alignment library (https://github.com/eseraygun/python-alignment). For the newly devised GLYcan SUbstitution Matrix (GLYSUM; Supplementary Table 4), the exhaustive list of *in silico* modifications resulting in glycans with observed glycowords was generated (n = 1,238,879). All thereby observed monosaccharide and/or bond substitutions were recorded in a symmetric matrix and converted into substitution frequencies by dividing them by the total number of retained modifications. The substitution score *S*_*ij*_ for each possible substitution was then calculated with the following formula:

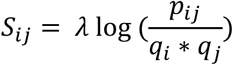

The substitution frequency is hereby denoted as *p*_*ij*_, while *q*_*i*_ and *q*_*j*_ describe the observed base frequencies of the respective glycoletters. Additionally, we used λ as a scaling factor (a value of four in this work) to arrive at suitable integer values by rounding all values up or down. Substitutions never observed during this procedure received a final value of -5, lower than any of the observed substitution scores, while the diagonal values of the substitution matrix were set at 5, higher than any of the observed substitution scores. The penalty for gaps for alignments in this work was set at -5, to match the minimal substitution score.

### Model training

All models were trained on an NVIDIA^®^ Tesla^®^ K80 GPU using PyTorch. Base model architecture and hyperparameters were optimized to minimize the averaged loss function across all taxonomic levels. A modified stratified shuffle split was used to randomly split glycans into training and validation sets so that, for every class, 80% of the glycans were present in the training set and 20% in the validation set. Further, only classes comprising at least five glycans were used for training and testing the model. We employed data augmentation by forming a generalizable subset of all possible isomorphic glycans, if a glycan sequence had isomorphic glycans. Specifically, we swapped the order of double branches and exhaustively exchanged the main branch with the side branches closest to the non-reducing end. The resulting sequence still described the same glycan in a slightly different way, increasing model robustness during training. Glycans were featurized into glycowords, brought to equal lengths using a padding token, and used in batches of 32 glycans for training and testing.

The final SweetOrigins models for each taxonomic level consisted of a three-layered, bidirectional recurrent neural network using long short-term memory (LSTM^43^) units with 128 nodes per layer, including an embedding layer for the glycowords and a fully connected layer at the end, predicting the taxonomic class of a glycan. The embedding layer was derived by first training a character-based model and then a glycoletter-based model, as described previously^14^. Then, the embedding was extracted and used to calculate initial glycoword embeddings for SweetOrigins. The last, fully connected layer was initialized by Xavier initialization^44^ and the number of nodes was determined by the number of unique taxonomic groups at the respective taxonomic level. We used a cross-entropy loss function and the ADAM optimizer with a starting learning rate of 0.0001 (decaying it with a cosine function over 100 epochs during training) and a weight decay of 0.005. Additionally, we employed an early stopping criterion after 10 epochs without improvement in validation loss for regularization.

### Phylogenetic tree construction

To construct phylogenetic trees from 16S rRNA, we retrieved 16S rRNA sequences from the corresponding species from the SILVA database^45^. We then used MAFFT^46^ (version 7.45) to construct a multiple sequence alignment with default parameters (except for using the scoring matrix 1PAM for closely related sequences) and a resulting phylogenetic tree using neighbor joining of conserved sites.

Phylogenetic trees from learned glycan embeddings were constructed by extracting the trained embedding layer of SweetOrigins to calculate glycan embeddings by averaging their glycoword embeddings. For each entry in a given taxonomic class, we generated a glycome embedding by averaging their glycan embeddings. From this, we calculated a cosine distance matrix and performed hierarchical clustering to construct dendrograms.

Analogously, we constructed phylogenetic trees from glycan alignments by first calculating a corresponding distance matrix. For this, we exhaustively aligned all possible pairs of glycans from two taxonomic classes. The average alignment score was then used as a measure of similarity. The resulting similarity matrix was converted into a distance matrix and normalized between zero and one. We built hierarchical clustering dendrograms based on this distance using the SciPy library with default parameters.

### Inferring the evolutionary origin for unlabeled glycans

For each unlabeled glycan sequence, glycans were aligned with all species-labeled glycans and ranked by alignment score. In parallel, the glycan sequence was used as input for all trained SweetOrigins models. The lowest taxonomic level resulting in concordance of the five highest scoring alignments and model inference was treated as the inferred origin of a glycan sequence. Briefly, if the species-level SweetOrigins and the best alignments concurred in their species prediction, the glycan was labeled with this species. If model and alignment disagreed, they were compared at the genus level, etc. If no agreement up to the domain level could be achieved, no taxonomic origin was inferred for the respective glycan.

### Data availability

Data used for all analyses can be found in the supplementary tables.

### Code availability

All code and trained models can be found at https://github.com/midas-wyss/sweetorigins.

## Acknowledgements

The authors would like to thank Jacqueline Valeri and Mathieu Groussin for helpful discussions.

## Contributions

D.B. conceived the method. D.B., D.M.C., and J.J.C. designed the experiments. D.B. performed the experiments and implemented the method. R.K.P. developed the GlycoBase web tool. D.M.C. and J.J.C. supervised the work. D.B., R.K.P., D.M.C., and J.J.C. wrote and approved the manuscript.

