## Supplementary Information for "SweetOrigins: Extracting Evolutionary Information from Glycans"

### Supplementary Figures

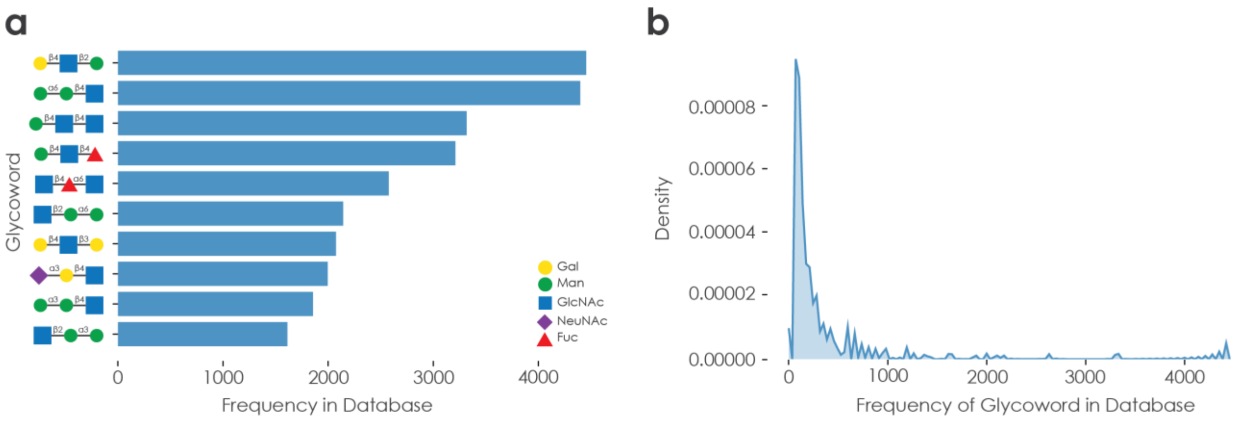


**Supplementary Fig. 1 | Glycoword distribution in the expanded glycan database.** The frequency of all 19,866 observed glycowords was determined in the database of 19,299 unique glycans. The 10 most frequent glycowords (**a**) as well as the kernel density estimate for all glycowords (**b**) are shown here. Glycans are drawn in accordance with the symbol nomenclature for glycans (SNFG).

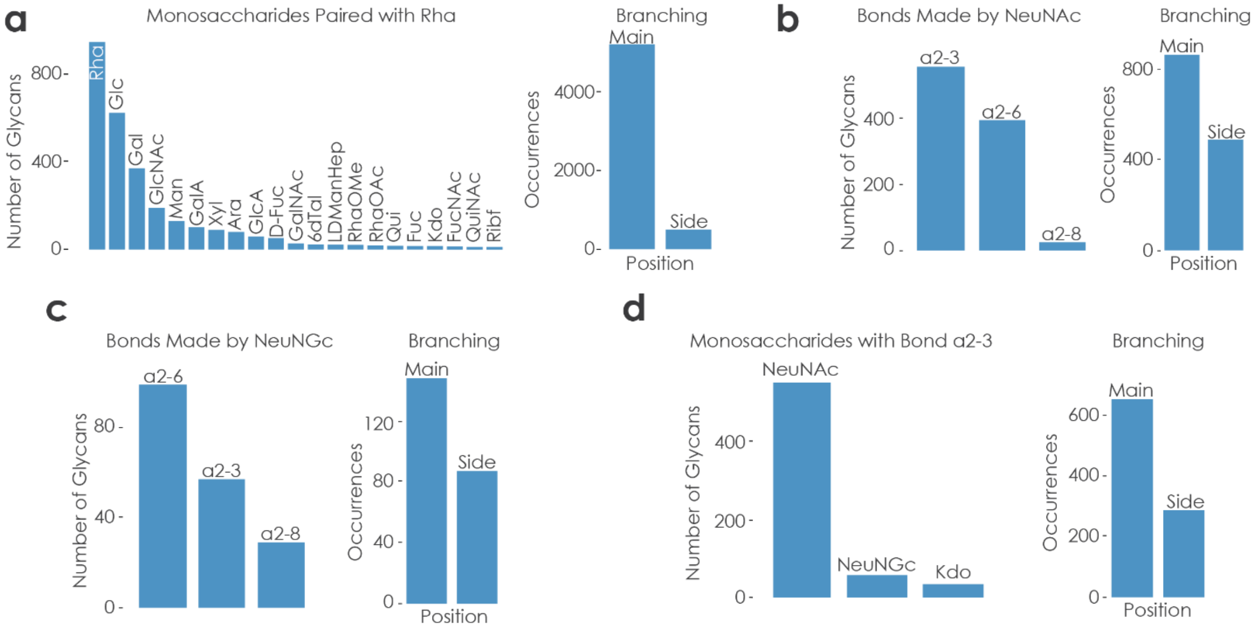


**Supplementary Fig. 2 | Typical local structural contexts of common monosaccharides and bonds.** The expanded glycan database presented here was used to investigate the typical binding partners of a range of glycoletters. Exemplary applications of this approach include the study of neighboring monosaccharides and branch position of the monosaccharide rhamnose (Rha, **a**), binding orientations and branch positions of N-acetylneuraminic acid (NeuNAc, **b**) as well as N-glycolylneuraminic acid (NeuNGc, **c**), and monosaccharides engaging in the binding orientation α2-3, as well as on which branch they are typically located (**d**). For visualization, we used a threshold of at least 10 co-occurrences in our database.

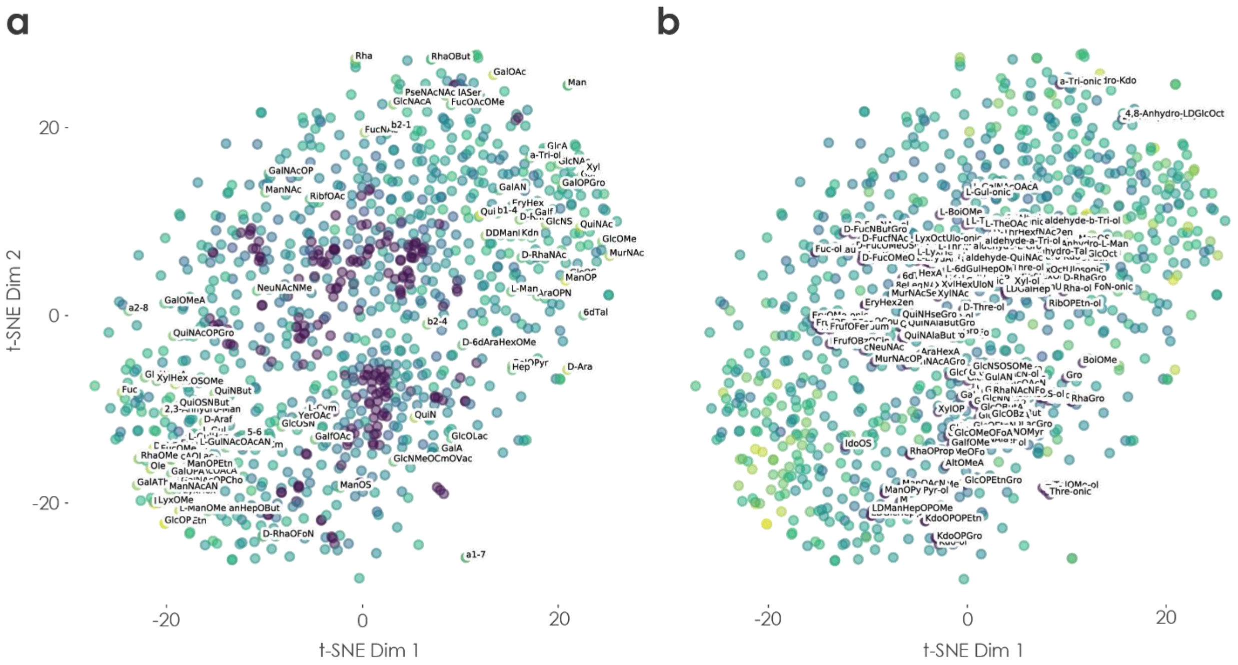


**Supplementary Fig. 3 | Changes in glycoletter representation after language model training.** After refining the pre-trained glycoletter embeddings via training a glycoletter-based language model, the Euclidean distance between pre-trained and trained embeddings was calculated for every glycoletter. Glycoletter embeddings exhibiting the most (more than 70% of the maximum change, **a**) as well as least (less than 10% of the maximum change, **b**) change during training are labeled. Glycoletter embeddings are colored from purple to yellow by Euclidean distance between pre-trained and trained embeddings. Glycoletters exhibiting large changes during training are characterized by higher frequency in our database, while the embedding of very rare glycoletters did not appreciably change due to lack of sufficient training examples.

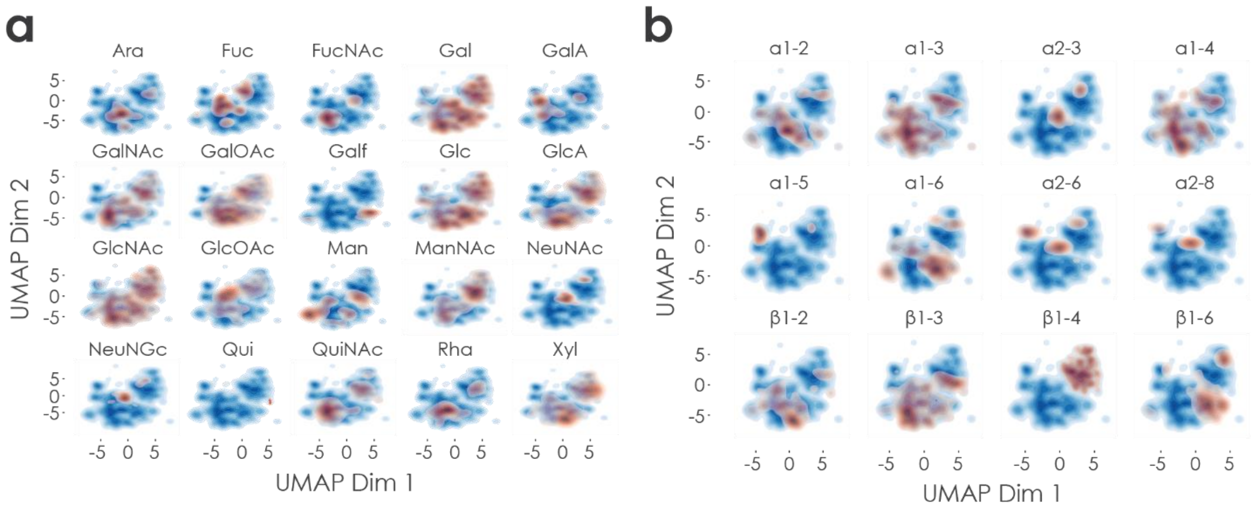


**Supplementary Fig. 4 | Trained glycoword embeddings colored by common monosaccharides and bonds.** For each observed glycoword, an embedding was constructed by averaging its constituent glycoletter embeddings. Kernel density estimate plots are shown for all glycowords (blue), colored red by the presence of one of a number of monosaccharides (**a**) or bonds (**b**) in the glycoword. Glycoword embeddings are visualized by dimensionality reduction via uniform manifold approximation and projection (UMAP).

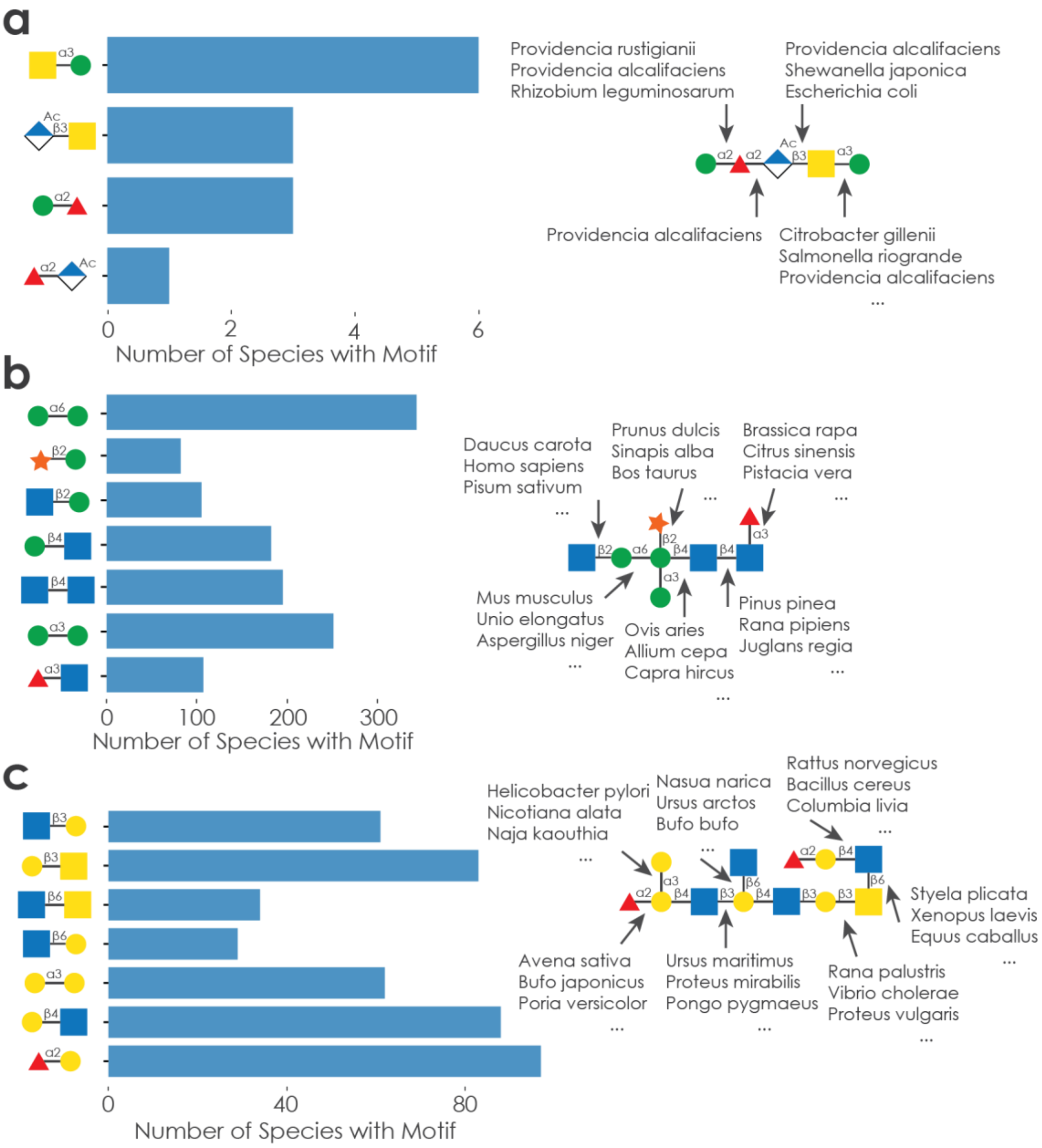


**Supplementary Fig. 5 | Generating biomining hypotheses for specific glycan linkages.** For several representative glycans (**a-c**), direct linkages between two monosaccharides were extracted and cross-referenced with all disaccharides observed in glycans of a given species for all species present in our database. For each disaccharide, the number of species exhibiting this motif is shown here. Additionally, up to three example species known to contain the respective disaccharide in at least one glycan are shown in the context of the glycan.

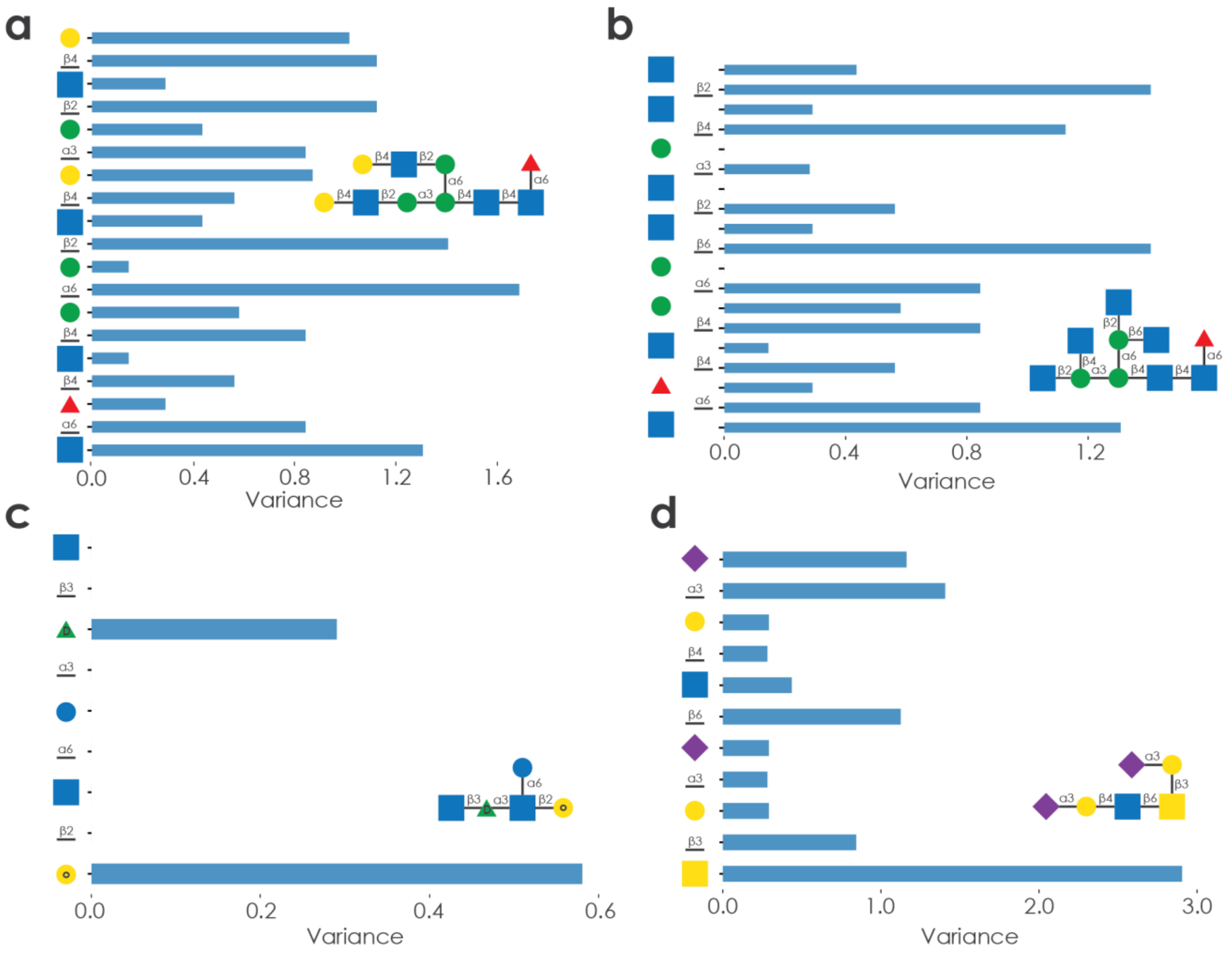


**Supplementary Fig. 6 | Sensitivity analyses of representative glycans at the glycoletter level.** For a set of representative glycans (**a-d**), exhaustive *in silico* modification was performed, only retaining modified glycans consisting of observed glycowords. For each position in a glycan sequence, the number of unique substitutions was then divided by the logarithm of the total number of monosaccharides or bonds observed in our database, respectively, to yield a measure of variance.

a

b

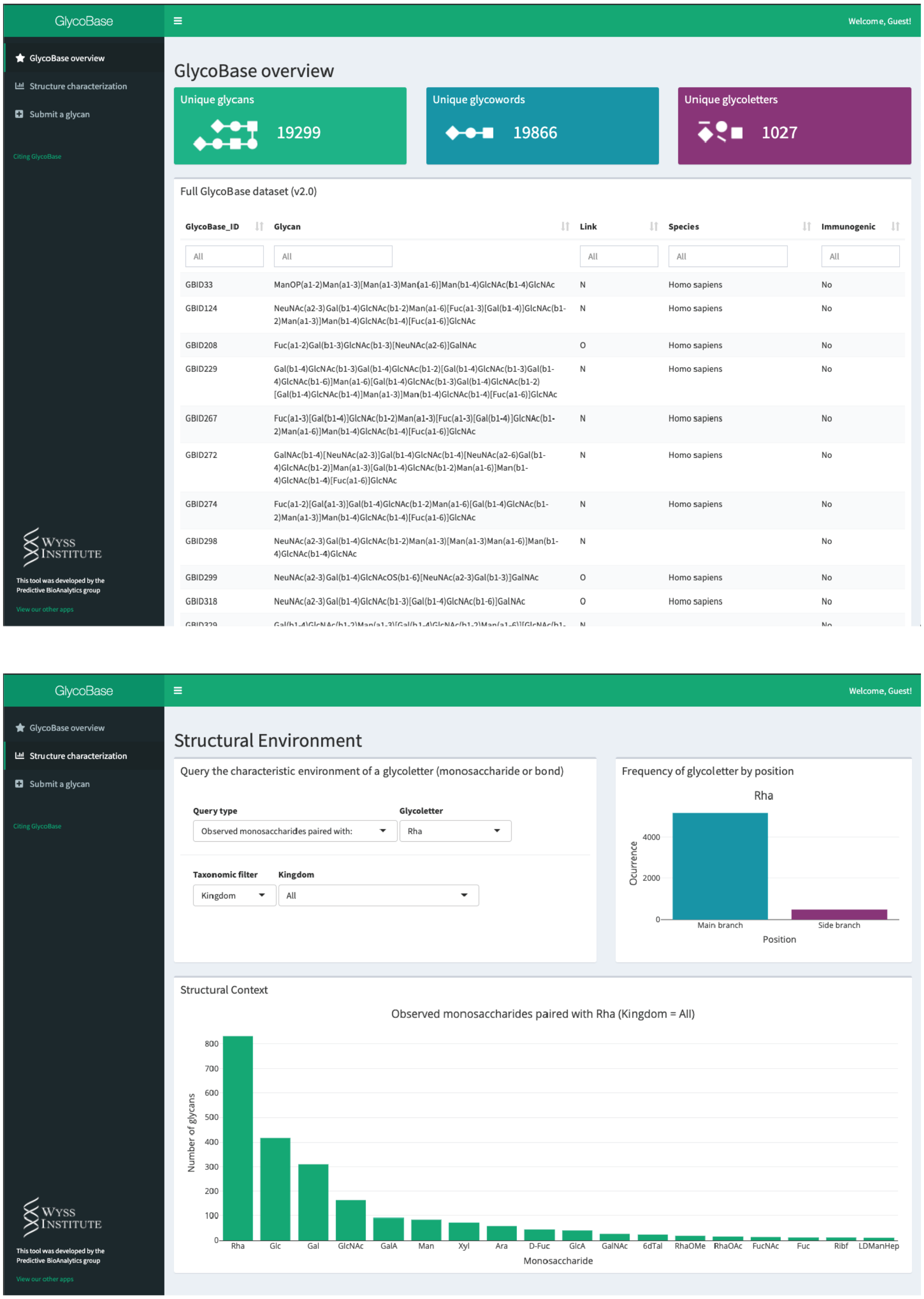


**Supplementary Fig. 7 | GlycoBase database interface.** Example screenshots of the GlycoBase main page (**a**) and the local structural context tool (**b**) are shown. Structural environment analysis of the monosaccharide rhamnose was performed with the herein developed GlycoBase platform. Next to the analysis of surrounding glycoletters and position of the queried glycoletter in the glycan, the analysis can also be restricted to a taxonomic kingdom of choice.

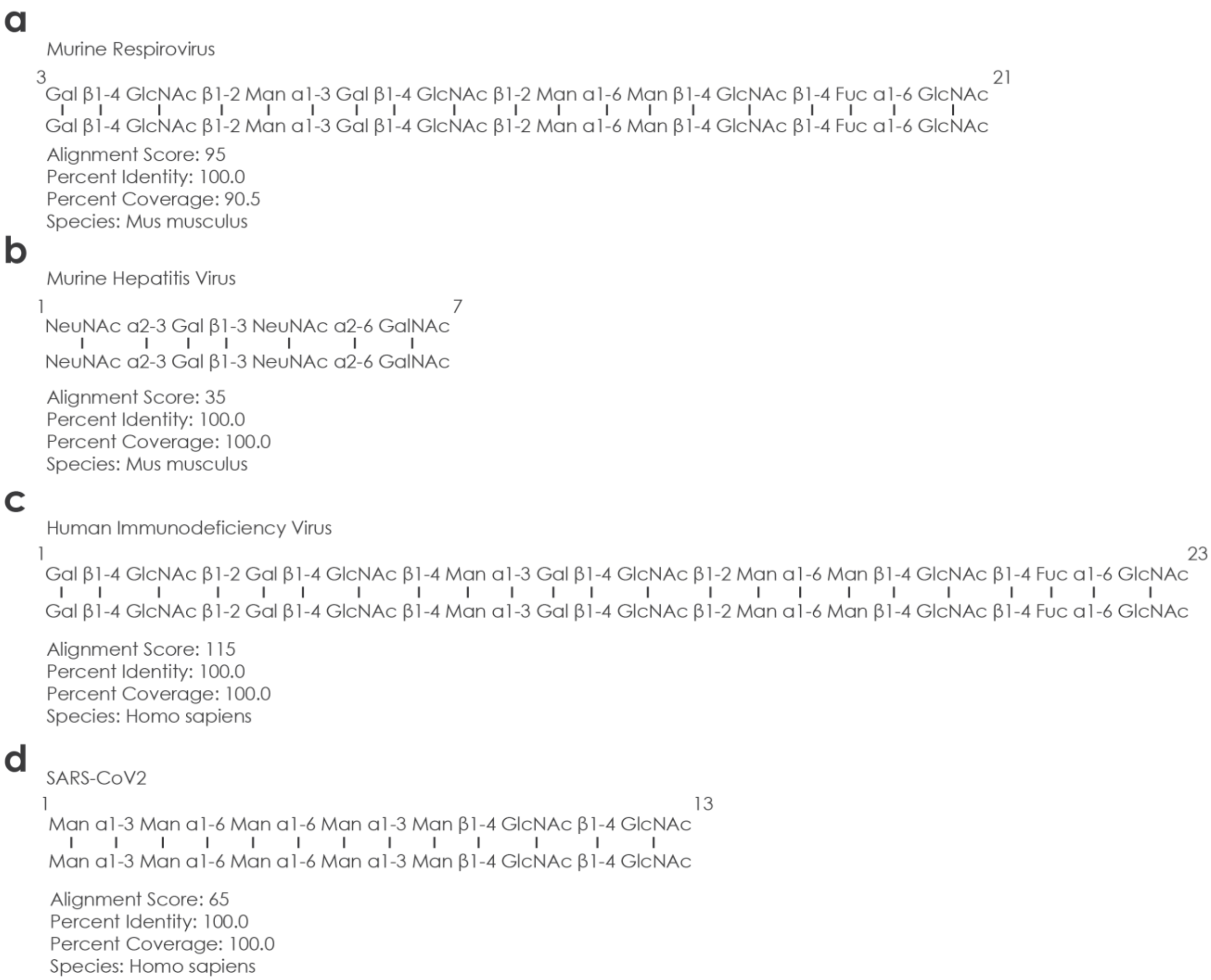


**Supplementary Fig. 8 | Viral glycans aligned to host glycans.** For representative viral glycans (**a-d**), we aligned their sequences to all glycans from their host organism as described in the methods section. The highest scoring alignment for each glycan is shown here, together with the associated alignment score, identity, coverage, as well as host species.

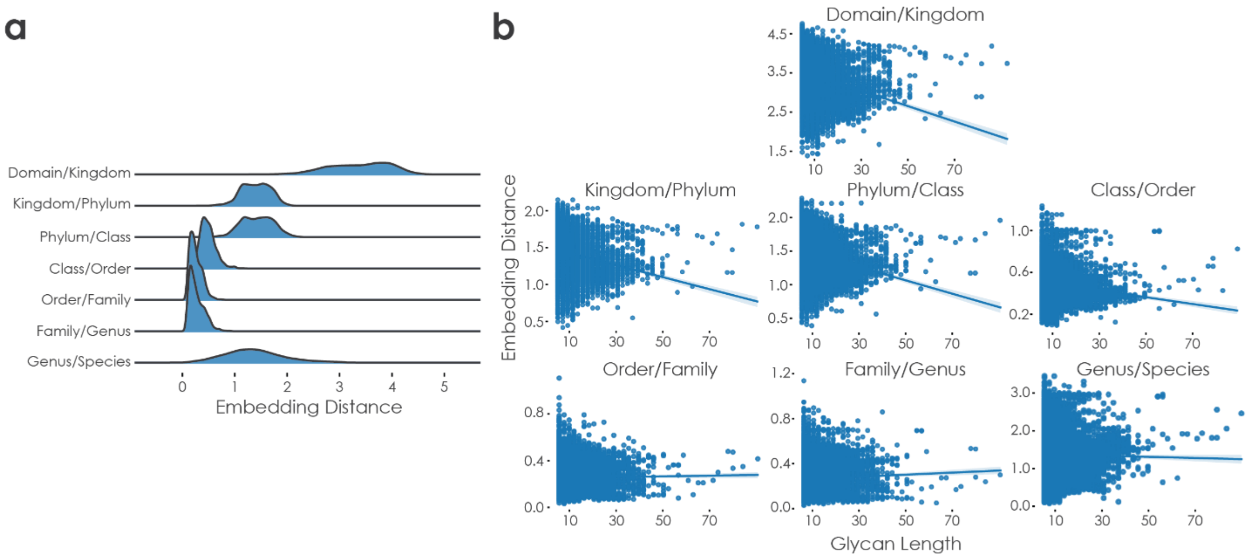


**Supplementary Fig. 9 | Embedding changes for SweetOrigins models trained on different taxonomic levels.** The embedding for all 8,966 unique glycans with a species label was formed from the trained embedding layers of SweetOrigins models for all taxonomic levels. Then, for each glycan, we calculated the Euclidean distance between embeddings of two subsequent taxonomic levels (e.g., the difference between embeddings of a domain-trained and a kingdom-trained SweetOrigins model). The distribution of embedding distances is shown as a ridgeline plot for each taxonomic comparison (**a**) and demonstrates large changes to the embedding for both coarse-grained (domain to kingdom, kingdom to phylum, phylum to class) as well as fine-grained taxonomic classifications (genus to species). Analyzing the dependence of embedding change on glycan length (**b**), fitted by a linear regression model, revealed that the pronounced embedding changes at the transitions from the domain, kingdom, and phylum level were mostly restricted to short glycans, potentially hinting at their limited expressiveness.

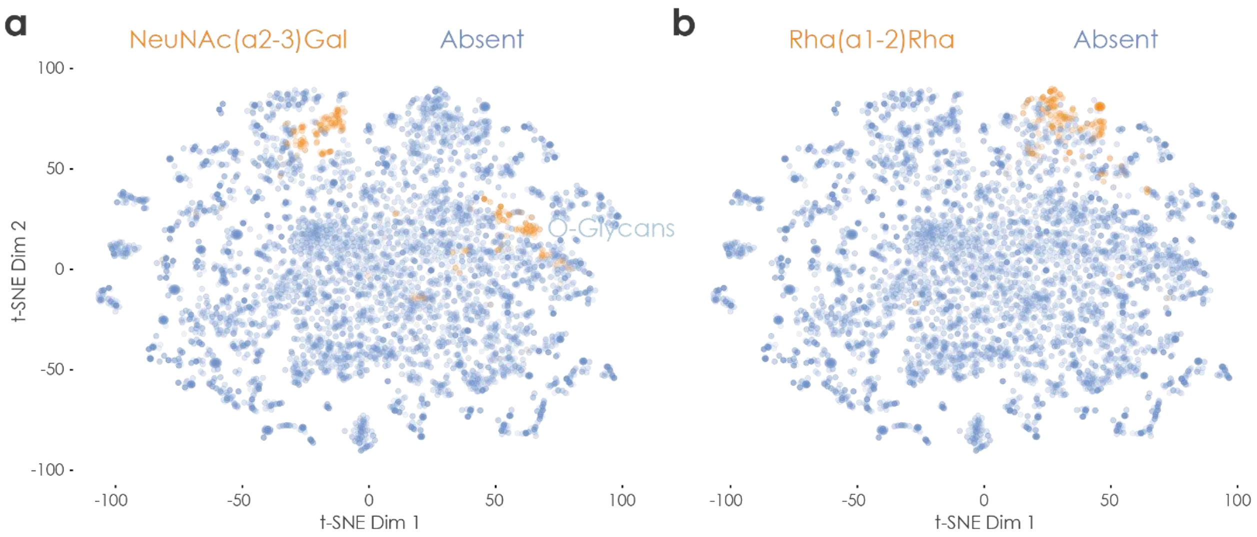


**Supplementary Fig. 10 | Tracking disaccharide motifs throughout evolutionary time.** All glycans with species information are shown in embedding space, plotted via t-SNE, analogous to Fig. 3d. For representative disaccharide motifs (**a-b**), all glycans containing the motif are colored in orange. Glycans containing NeuNAc(α2-3)Gal are enriched for N-/O-linked glycans of vertebrates as well as bacterial glycans of the phyla Bacteroidetes, Firmicutes, and Proteobacteria (**a**), while glycans containing Rha(α1-2)Rha are restricted to a region mostly containing Proteobacteria, Angiosperms, and Ascomycota (**b**).

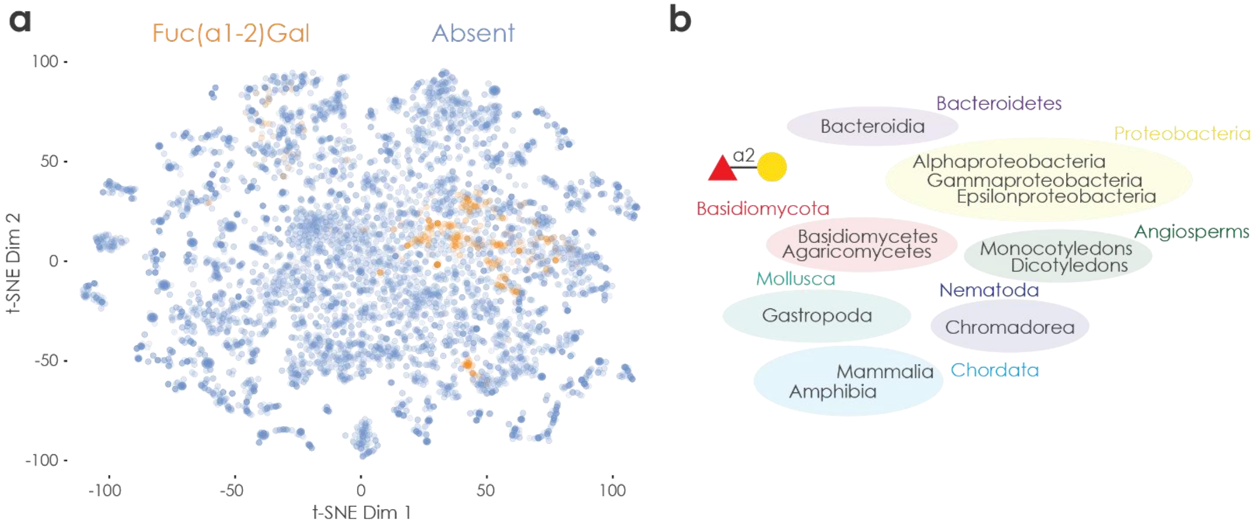


**Supplementary Fig. 11 | Evolutionary tracking of Fuc(α1-2)Gal. (a**) All glycans with species information are shown in embedding space, plotted via t-SNE, analogous to Fig. 3d. Glycans containing the motif Fuc(α1-2)Gal are colored in orange. (**b**) All taxonomic classes that contained glycans with the shown motif are depicted and colored by their phyla.

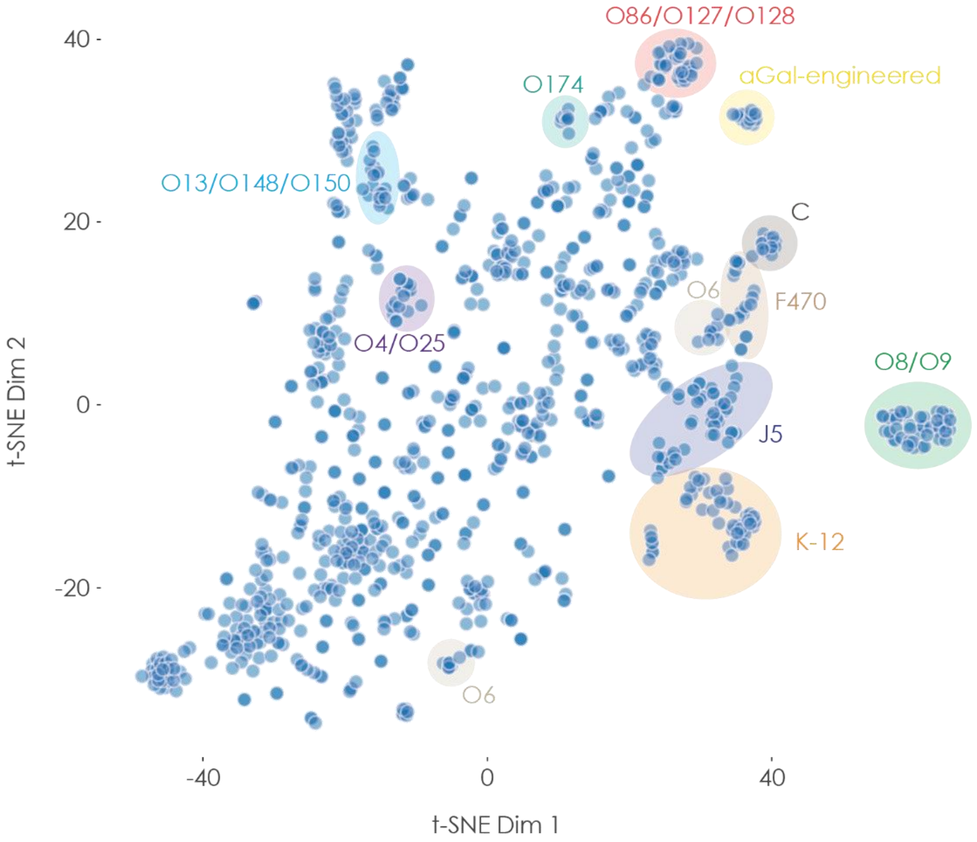


**Supplementary Fig. 12 | Glycans of *Escherichia coli* in embedding space distinguish strains.** We calculated the embedding for all 1,010 *E. coli*-derived glycans with strain information with the embedding layer of the trained species-level SweetOrigins model, plotted them via t-SNE, and colored areas that are enriched for annotated *E. coli* strains.

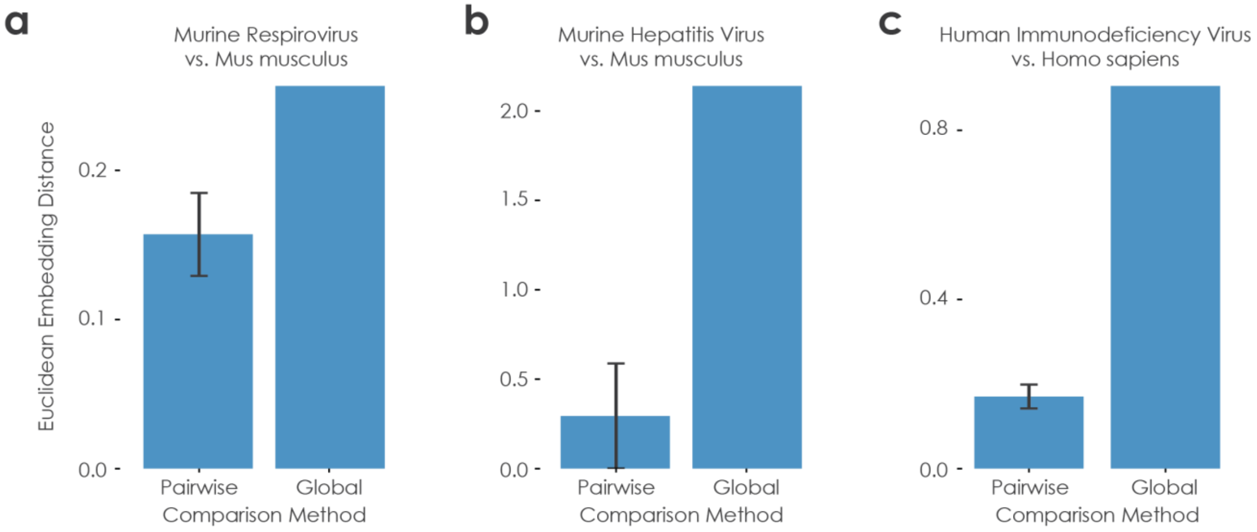


**Supplementary Fig. 13 | Embedding distance between viral glycans and those of their host species.** For representative viruses (**a-c**), the distance between viral glycans and glycans of their host was determined by two routes. For pairwise distances, each viral glycan was aligned to all host glycans and the Euclidean distance between the embeddings of the viral glycan and the host glycan with the highest alignment score was calculated. Shown is the mean pairwise distance between viral and host glycans ± s.e.m. For the global distance, the embeddings of all glycans of a given virus were averaged and the Euclidean distance with regard to the average embedding of all host glycans was calculated and is shown here. While individual viral glycans are very similar to identical compared to host glycans, viral glycans only comprise a subset of host glycans and therefore differ more on a global level.

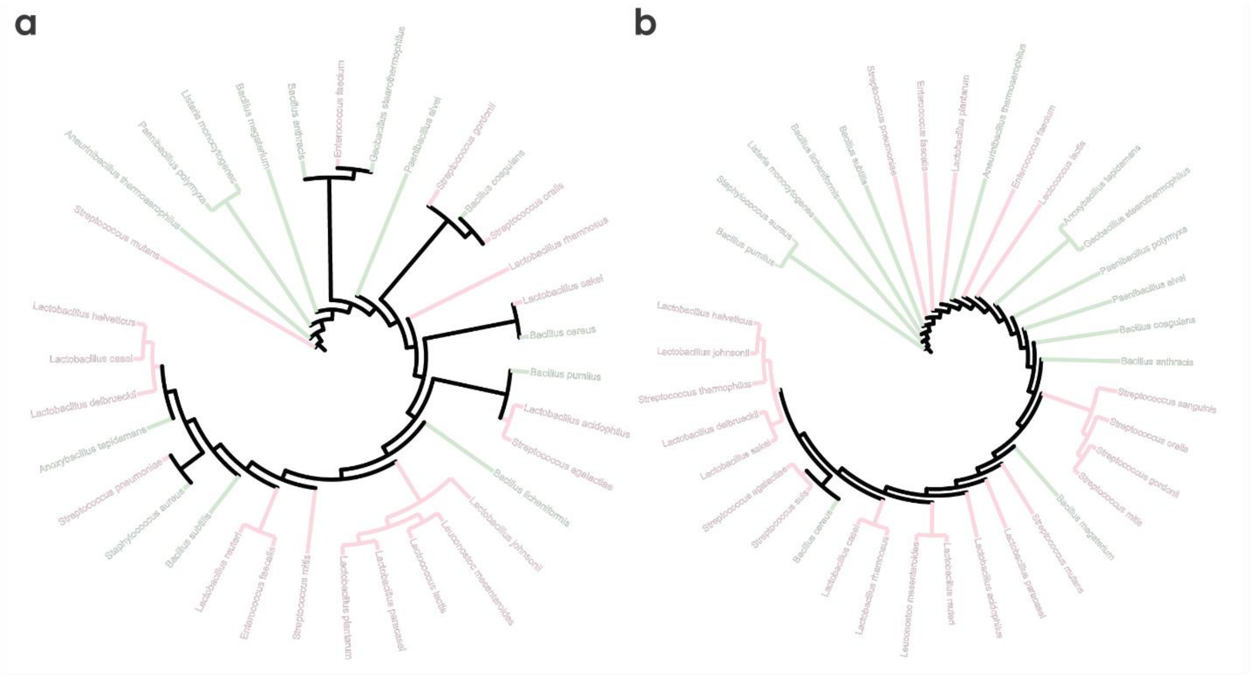


**Supplementary Fig. 14 | Phylogenetic tree of Bacilli based on glycan embeddings and alignments.** For all species in the taxonomic class Bacilli (phylum Firmicutes) with at least five glycans in our dataset, we calculated their glycan embedding from the trained species-level SweetOrigins model and averaged them by species (**a**) or calculated their exhaustive pairwise glycan alignment score (**b**). Then, we calculated their pairwise distances and performed hierarchical clustering, yielding a dendrogram visualized by Interactive Tree Of Life v5.5. Species were colored according to their taxonomic order (pink Lactobacillales, green Bacillales).

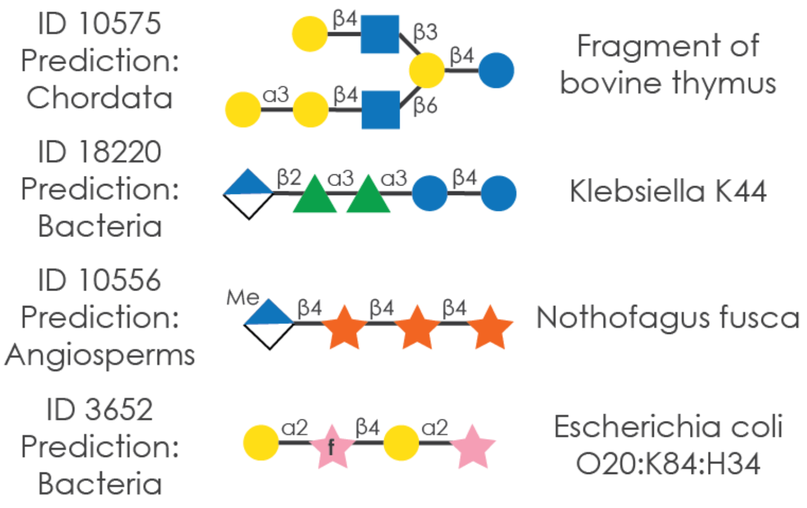


**Supplementary Fig. 15 | Validated predictions of evolutionary origins of glycans.** Origin inferences made for representative glycans were validated by targeted literature searches. For each glycan, the predicted taxonomic class as well as the actual species a glycan originated from is shown. Glycans are drawn in accordance with the symbol nomenclature for glycans (SNFG).
